# *Photorhabdus* Dam methyltransferase overexpression impairs virulence of the nemato-bacterial complex in insects

**DOI:** 10.1101/545061

**Authors:** Amaury Payelleville, Dana Blackburn, Anne Lanois, Sylvie Pagès, Marine Cambon, Nadège Ginibre, David Clarke, Alain Givaudan, Julien Brillard

## Abstract

*Photorhabdus luminescens* is an entomopathogenic bacterium found in symbiosis with the nematode *Heterorhabditis*. Dam DNA methylation is involved in the pathogenicity of many bacteria, including *P. luminescens,* whereas studies about the role of bacterial DNA methylation during symbiosis are scarce. The aim of this study was to determine the role of Dam DNA methylation in *P. luminescens* symbiosis with *H. bacteriophora*. We constructed a strain overexpressing *dam* by inserting an additional copy of the *dam* gene under the control of a constitutive promoter in the chromosome of *P. luminescens* and then achieved association between this recombinant strain and nematodes. The *dam* overexpressing strain was able to feed the nematode *in vitro* and *in vivo* similarly as a control strain, and to re-associate with Infective Juvenile (IJ) stages in the insect. No difference in the amount of emerging IJs from the cadaver was observed between the two strains. Compared to the nematode in symbiosis with the control strain, a significant increase in LT_50_ was observed during insect infestation with the nematode associated with the *dam* overexpressing strain. These results suggest that the *P. luminescens* Dam plays a role in the pathogenicity of the nemato-bacterial complex.

## Introduction

Studies aiming to understand bacteria-host interactions often show that molecular mechanisms involved in mutualism or pathogenesis are shared [1]. This raises the interest to study models that have a life-cycle including both mutualism and pathogenicity stages. *Photorhabdus luminescens* (*Enterobacteriaceae*) is symbiotically associated with a soil nematode, *Heterorhabditis bacteriophora* [2]. The nemato-bacterial complexes are highly pathogenic for insects and used as biocontrol agents against insect pest crops [3]. Mutualistic interaction between both partners is required as *Photorhabdus* is not viable alone in the soils and *Heterorhabditis* cannot infect and reproduce without its symbiont [4]. *Photorhabdus* is carried inside the nematode gut during the infective juvenile stage (IJ), a stage that is similar to the well characterized dauer-stage of *Caenorhabditis elegans* [5]. After their entrance by natural orifices such as stigmata, or by cuticle disruption, nematodes release *Photorhabdus* in the hemocœl of the insect [6, 7]. The bacteria then grow and produce a broad-range of virulence factors to kill the insect by septicemia within 48 to 72 hours [8, 9]. Regurgitation and multiplication of the symbiont induce a phenomenon called “IJ recovery” resulting in the formation of a self-fertile adult hermaphrodite from every IJ [7]. Nematodes feed specifically on their symbiotic bacteria [10, 11]. Once nutrients are lacking and nematodes have done several development cycles, some bacterial cells adhere to hermaphrodite gut at INT9 cells [12]. Bacteria which can adhere to these cells express the Mad pilus [12, 13]. Hermaphrodites lay about 100 to 300 eggs giving rise to IJs feeding on and re-associating with *Photorhabdus*. Some eggs are not released and develop inside the hermaphrodite by a mechanism called *endotokia matricida* [14]. Nematodes coming from *endotokia matricida* will become IJs only and will re-associate with *Photorhabdus* inside the hermaphrodite [14, 15]. After re-association of both partners, the complexes exit from the cadaver to reach the soil in order to infect other insects [16]. The pathogenic cycle implies a strong interaction between the bacterium and the nematode and requires a bacterial switch from mutualism to pathogenic state. It is therefore a good model to study differences between both states [17].

In enterobacteria, Dam (for DNA Adenine Methyltransferase) adds an m6A methylation mark to the adenine of 5’-GATC-3’ sites. It can be involved in epigenetic mechanisms because of a binding competition between a transcriptional regulator and Dam for some promoter regions, leading to differential gene transcription [18]. Dam DNA methylation plays a role in the pathogenicity of several pathogens such as *S.* Typhimurium [19, 20], *Y. pestis* and *Y. pseudotuberculosis* [21, 22]. Other DNA methylation marks (m4C and m5C) involved in pathogenicity such as in *H. pylori* [23, 24] have also been described. However, the involvement of DNA methylation in mutualistic associations are focused on host modifications, whereas bacterial DNA methylation data are scarce and limited to bacterial-plant interactions [25-27]. Recently we showed that the overexpression of *dam* in *P. luminescens* decreases motility and virulence and increases biofilm formation [28]. Here, we focused on the symbiotic stages of *P. luminescens* life-cycle. We constructed a strain overexpressing Dam MTase with a chromosomal insertion and achieved a symbiosis between this strain and the nematode *H. bacteriophora*. The involvement of Dam in symbiosis was studied after insect infection with the nemato-bacterial complex. The insect mortality rate over time, the IJs emergence from the cadaver and the number of bacteria associated with these IJs were quantified.

## Material and methods

### Strains, plasmids and growth conditions

The bacterial strains, nematode strains and plasmids used are listed in Table 1. Bacteria were grown in Luria broth (LB) medium with shaking at 28 °C for *Photorhabdus* and 37 °C for *E. coli*, unless stated otherwise. When required, IPTG was added at 0.2 mM, pyruvate at 0.1 % and sucrose at 3 %, antibiotics were used: gentamycin (Gm) at 20 μg/mL^-1^and chloramphenicol (Cm) at 8 μg/mL^-1^. Phenotypic characterization of the strains was determined as previously described [28].

**Table 1:**
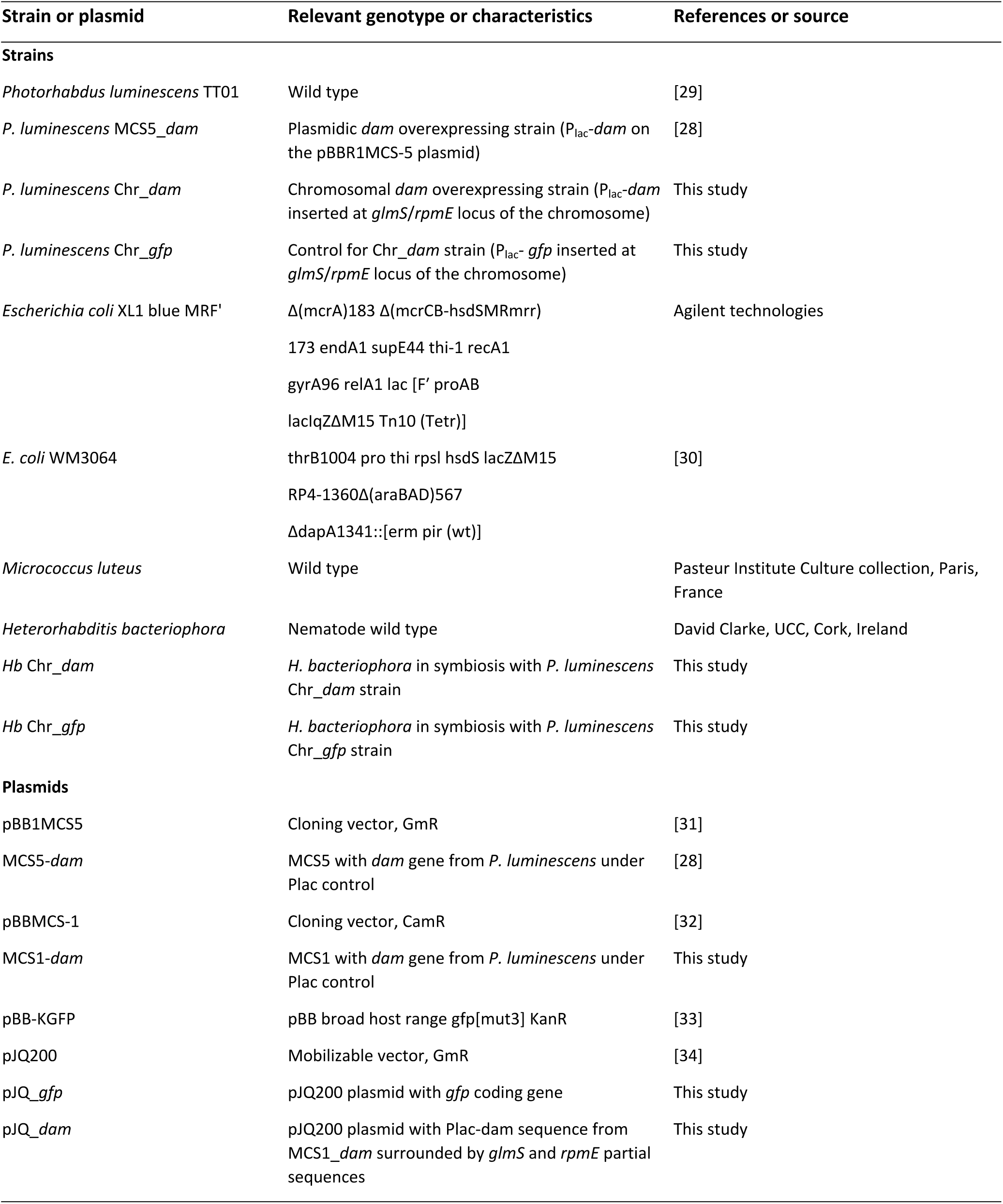
Strains and plasmids used in this study

### Chromosomal integration of *dam*

To avoid studying the effect of Dam overexpression on the bacterial nematode association using an instable plasmid-borne *dam* construction, we inserted the *dam* gene under the control of the promoter P_*lac*_ at the *rpmE/glmS* intergenic region of the chromosome [35] as follows. The *dam* gene was extracted from MCS5_*dam* plasmid [28], digested with *SalI* and *XbaI* enzymes (NEB) and the resulting 889 bp fragment was cloned in the pBB-MCS1 vector using T4 DNA Ligase (Promega). This plasmid MCS1_*dam* was then digested with *AatII* and *SacI* enzymes to obtain a DNA fragment of 2194 bp containing a chloramphenicol resistance gene and the *dam* gene controlled by the P_*lac*_ promoter. In parallel, a 643 bp fragment overlapping *glmS* gene and a 752 bp fragment overlapping *rpmE* gene from *Photorhabdus* were amplified using R_GlmS_SalI, F_GlmS_AatII and R_RpmE_SacI, F_RpmE_SpeI respectively (Table S1) and digested with the appropriate enzymes. Finally, the pJQ200 plasmid (Table 1) was digested by *SalI* and *SpeI* and ligated together with the three fragments. *E. coli* XL1 Blue MRF’ was transformed with the pJQ_Cam_P_lac-_*dam* ligation mixture and clones with the appropriate antibiotic resistance (i.e., CmR and GmR) were selected. Similarly, the pJQ_Cam_P_lac-_*gfp* plasmid was constructed using *gfp-mut3* gene (*Kpn*I-*Pst*I fragment) from pBB-KGFP (Table 1) instead of *dam.* The plasmid constructions were controlled by sequencing of the inserts.

The recombinant plasmids pJQ_Cam_P_lac-_*dam* or pJQ_Cam_P_lac-_*gfp* were then transferred in *P.luminescens* by conjugation as previously described [28]. The transconjugants were selected with both Cm and Gm. The allelic exchanges were performed on at least 20 independent transconjugants as previously described [36]. Finally, Sac resistant, Cm resistant and Gm sensitive clones were grown overnight in LB + Cm. Genomic DNA was extracted using QIAamp DNA Mini kit (Qiagen) and correct insertion was verified by sequencing the PCR fragment overlapping the insertion site (using primers L_verif_GlmS and R_verif_RpmJ). Clones with the correct insertion (Chr_*dam* and Chr_*gfp*) were then tested for their phenotypes as previously described [28] and conserved in glycerol (Table S2).

### RT-qPCR analysis

To quantify the level of *dam* overexpression in the Chr_*dam* strain, quantitative reverse transcription-PCR (RT-qPCR) were performed as previously described [28, 37]. Briefly, RNA samples from 3 independent cultures for each strain (Chr_*dam* and Chr_*gfp*) were extracted with RNeasy miniprep kit (Qiagen). Primers used are listed in Table S1. Results are presented as a ratio with respect to the housekeeping gene *gyrB*, as previously described [38].

### Insect virulence assay

*P. luminescens* Chr_*dam* and Chr_*gfp* strains virulence were tested for their virulence properties on *Spodoptera littoralis* in three independent experiments, as previously described [36]. Briefly, 20 µL of exponentially growing bacteria (DO_540nm_ = 0.3) diluted in LB, corresponding to about 10^4^ CFU for each strain were injected into the hemolymph of 30 sixth-instar larvae of *S. littoralis* reared on an artificial diet [39] with a photoperiod of L16:D8. Each larva was then individually incubated at 23 °C and mortality times were checked. Survival rate for each bacterial strain infestation were then analyzed with Wilcoxon test performed as previously described [36, 40] using SPSS V18.0 (SPSS, Inc., Chicago, IL) to compare the time needed to kill 50 % of the infested larvae.

### Nemato-bacterial monoxenic symbiosis

A nemato-bacterial complex between *H. bacteriophora* and *P. luminescens* Chr_*dam* or Chr_*gfp* strains was generated as follows. *Photorhabdus* WT strain was grown overnight at 27 °C with shaking in LB + pyruvate, plated on lipid agar plates [41] and then incubated at 27 °C during 48 h. 5000 IJs were added to *Photorhabdus* lipid agar plates and incubated during 4 days at 27 °C. Hermaphrodites were collected from lipid agar plates in 50 mL conical tubes by adding PBS to the plate, swirling and dumping into the tube. After hermaphrodites have settled, PBS was removed. This step was repeated until a clear solution was obtained. Egg isolation from hermaphrodites was then performed as follows. 200 μL of washed hermaphrodites were put into 3.5 mL of PBS. 0.5 mL of 5M NaOH mixed with 1mL of 5.6 % sodium hypochlorite was added and the tube was incubated for 10 minutes at room temperature with short vortex steps every 2 minutes. The tube was centrifuged (30 s, 1300 g) and most of the supernatant was removed leaving 100 μL in the tube. PBS was then added to a final volume of 5 mL. After vortexing and centrifugation, eggs were washed again with 5 mL PBS and collected after another centrifugation step. *P. luminescens* Chr_*dam* and the control strain were grown in 5 mL of LB overnight at 27 °C with shaking. 30 μL of the culture were spread on split lipid agar plates and incubated at 27 °C for two days prior to harvesting eggs. Equal amounts of eggs (~1000) were added to each plate. PBS was added to the empty part of the plate and plates were incubated for two weeks at 27 °C. IJs were collected in the PBS side of the plate and stored at 4 °C.

### Insects infestation and IJs emergence

*G. mellonella* infestations were performed in 1.5 mL Eppendorf tube to inhibit their weave ability that occurs in plates and which would hinder direct contact with EPN. In each tube, 100 μL of PBS containing 50 IJs were added on a filter paper and one *Galleria* larva was added. Tubes were incubated at 23 °C. *S. littoralis* infestations were performed in 12 well plates using filter papers containing 50 IJs as described above. One *S. littoralis* larva was added in each well with artificial diet. For both insects infestation, mortality was checked regularly over time during 72 hours. The survival rates for each nemato-bacterial complex were analyzed with Wilcoxon test performed as previously described [36, 40] using SPSS V18.0 (SPSS, Inc., Chicago, IL) to compare LT_50_ of the infested larvae.

### Bacterial CFUs in nemato-bacterial complex

CFUs for each nemato-bacterial complex were quantified as follows. IJs were filtered using a 20 μm pore-size filter to remove bacteria present in the solution. After resuspension in 5 mL of PBS, two additional PBS washing steps were performed. Then, 10 IJs were counted under binocular magnifier and placed in 10-50 μL volume in 1.5 mL tube. Manual crushing was performed using plastic putter and efficiency of nematodes disruption was verified by microscope observation. After addition of 1 mL LB, 100 µL of the suspension was plated on LB Petri dish, pure or at 10^-1^ dilution, with 3 replicates for each dilution. *Photorhabdus* CFUs were determined using a Li-Cor Odyssey imager and Image Studio version 1.1.7 version to discriminate luminescent colonies (corresponding to *P. luminescens*) from others. For each stain, three independent cultures were used to infect 3 insects, for a total of nine infestations. To test for differences in bacterial retention of IJs obtained from these infestations, we performed a generalized linear mixed model (glmm) including the identity of the strain culture as a random effect, using the spaMM package [42].

### Ethics statement

According to the EU directive 2010/63, this study reporting animal research is exempt from ethical approval because experiments were performed on invertebrates animals (insects).

## Results

### Effect of *dam* overexpression by chromosomal insertion on *P. luminescens* phenotypes

*dam* expression was quantified in the Chr_*dam* strain harboring an additional copy of the *dam* gene under the control of a strong promoter by a chromosomal insertion. An increase of 14-fold changes in *dam* expression in the Chr_*dam* strain was observed (p-value = 0.001, Confidence Interval 95% = 5,785 - 41,381) compared to the control strain Chr_*gfp* (harboring a *gfp* gene inserted on the chromosome).

To determine if the *dam* overexpression modified some *P. luminescens* phenotypes, similarly as a strain overexpressing *dam* using a plasmid did [28], we compared motility and insect pathogenicity of Chr_*dam* and Chr_*gfp* strains (control). A significant decrease in motility was observed for the Chr_*dam* strain (p-value < 10^-3^, Wilcoxon test) at 36h hours post inoculation (Fig 1A). LT_50_ in *S. littoralis* was significantly reduced (p-value < 10^-3^, Wilcoxon test) in the *dam* overexpressing strain compared to the control strain, with a delay of 2 hours (32.8 hours for the control and 34.9 for Chr_*dam* strain; Fig 1B). These data confirmed that the *dam* overexpression in *P. luminescens* impairs the bacterial virulence in insect. No other tested phenotype was impacted by chromosomal *dam* overexpression in *P. luminescens* (Table S2).

**Figure 1:**
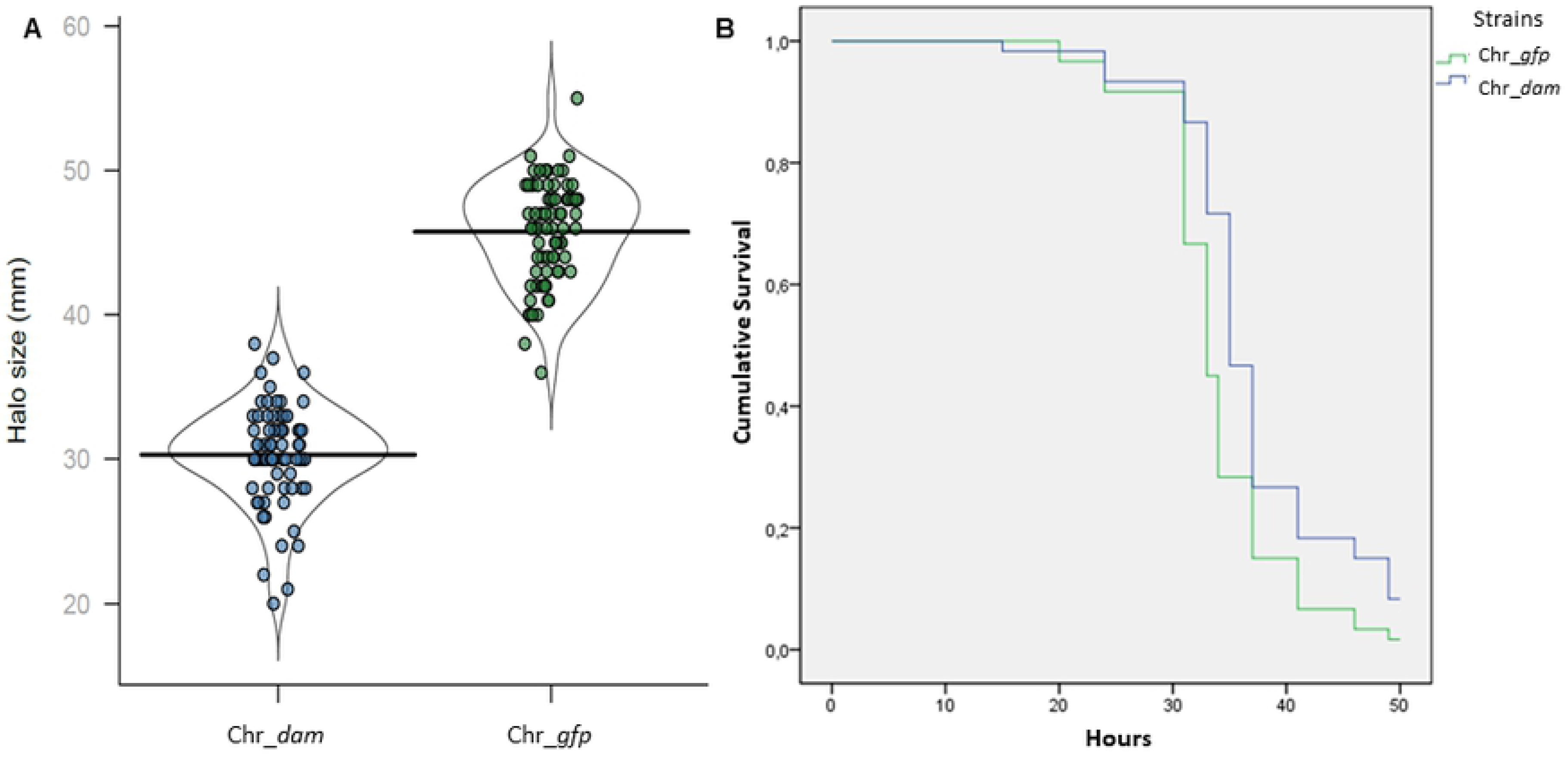
Motility and pathogenicity of Chr_*dam* strain. (A) Violin-plot of motility halo size for Chr_*dam* and Chr_*gfp* strain after 36 hours of growth on motility medium. The difference between the two strains was significant (Wilcoxon test, p-value<0.001). (B) Survival of *S. littoralis* larvae after injection of 10^4^CFU of Chr_*gfp* (green) or Chr_*dam* (blue). Chr_*dam* strain was significantly delayed (2hours) in the time needed to kill 50 % of the larvae (Wilcoxon test, p-value<0.001).

### Symbiosis establishment

To study Dam involvement in the symbiosis stage of *P. luminescens* life-cycle, the construction of a complex between *P. luminescens* Chr_*dam* or Chr_*gfp* strains and *Heterhorhabditis* was performed. No difference in the number of emerging IJs *in vitro* could be detected for the three biological replicates (Fig S1). This suggests that the nematode can feed and establish a symbiotic relationship with the Chr_*dam* strain in *in vitro* conditions.

### Pathogenicity of the EPN complex in *G. mellonella* and *S. littoralis*

In order to study the role of the *P. luminescens* Dam MTase in the virulent stage of the nemato-bacterial complex, *G. mellonella* or *S. littoralis* were infested and insect larvae mortality was monitored overtime. Both nemato-bacterial complexes (i.e., nematodes in symbiosis with either Chr_*dam* or Chr_*gfp* strains, respectively *Hb* Chr_*dam* and *Hb* Chr_*gfp*) were pathogenic as they caused insect death in less than 72 hours. For *G. mellonella*, the LT_50_ were 48 and 50.6 hours for *Hb* Chr_*gfp* and *Hb* Chr_*dam*, respectively. The difference between the two strains was significant (p-value<0.05, Wilcoxon test) (Fig 2A). In *S. littoralis* the LT_50_ was delayed by almost 6 hours (48.4h and 54.2h for *Hb* Chr_*gfp* and *Hb* Chr_*dam*, respectively) (Fig 2B). This difference was highly significant (p-value <0.001, Wilcoxon test).

**Figure 2:**
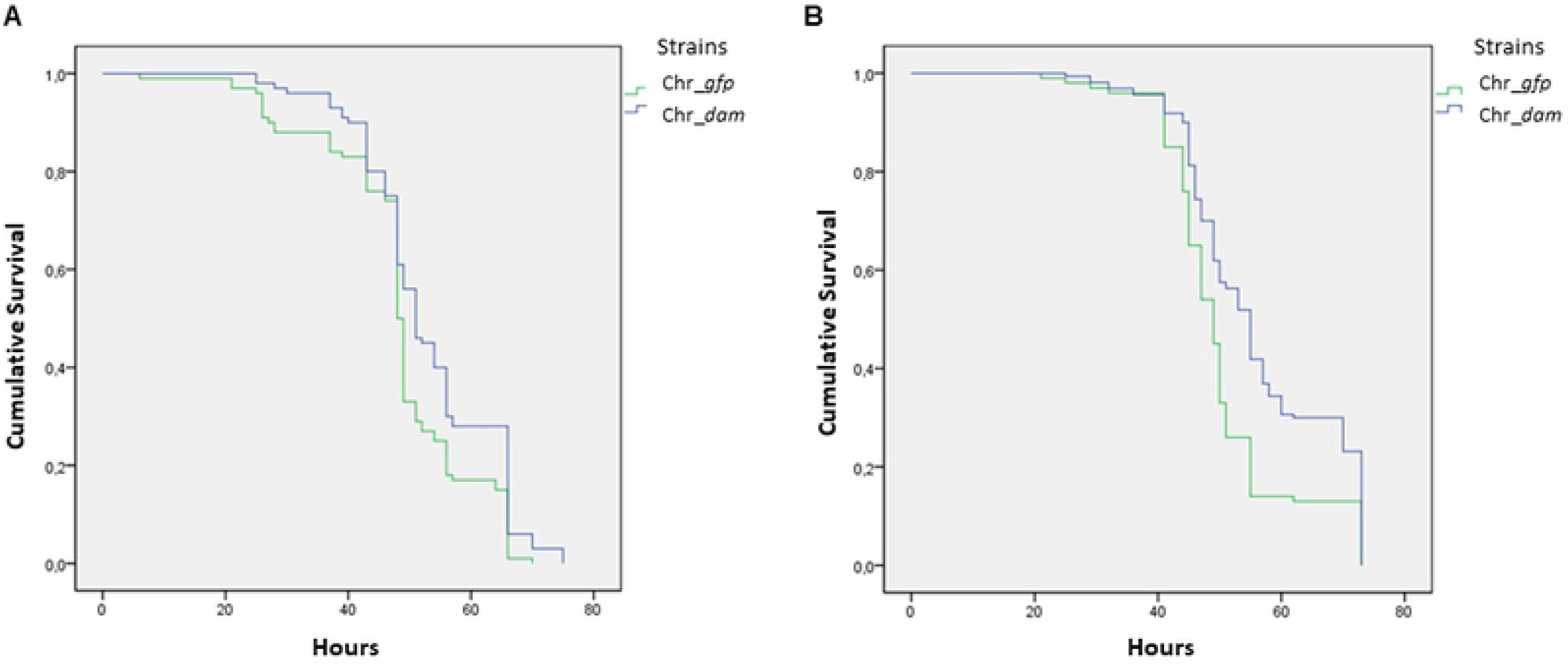
Nemato-bacterial complex pathogenicity by infestation. (A) Survival of *G. mellonella* larvae after infestation by 10 nematodes associated with Chr_*gfp* bacterial strain (green) or Chr_*dam* strain (blue). A significant difference of 2 hours was observed for the time needed to kill 50 % of the larvae between the two strains (Wilcoxon, p-value<0.05). (B) Survival of *S. littoralis* larvae after infestation as described above. A significant difference was observed with an almost 6 hours delay for the Chr_*dam* strain (Wilcoxon, p-value<0.001).

### Emerging IJs from cadavers

To investigate Dam role in the *in vivo* association between the nematode and *P. luminescens,* we quantified IJs emerging from each insect larvae. The amount of emerging IJs exiting from the cadavers of *G. mellonella* and *S. littoralis* were not different between both nemato-bacterial complexes used (p-value = 0.991 and p-value = 0.31, respectively, Wilcoxon test) (Fig 3A and 3B).

**Figure 3:**
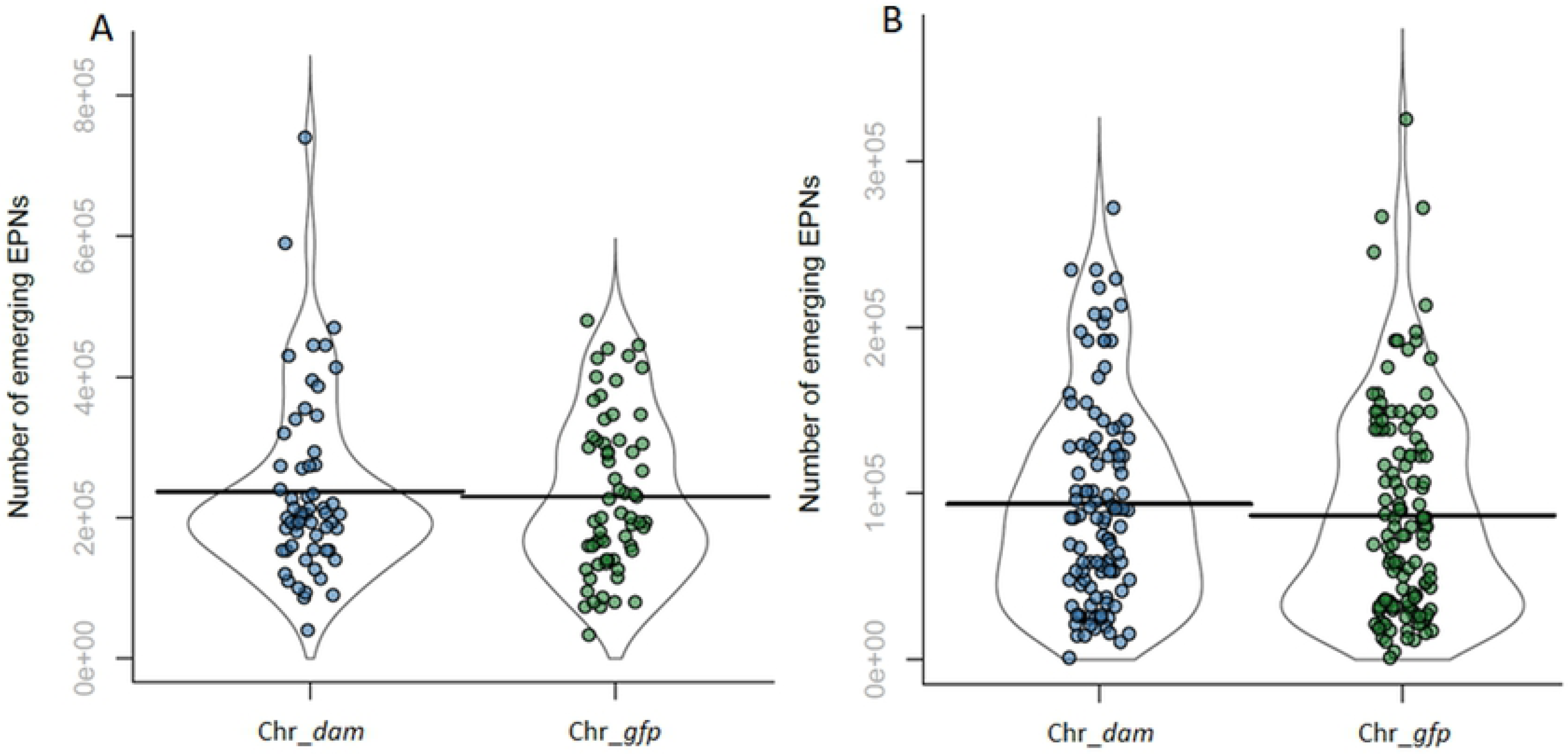
Number of emerging IJs from each cadaver. (A) Emerging IJs from each *G. mellonella* cadaver for each strain. The amount of IJs exiting from larvae cadaver were not significantly different between the two strains (Wilcoxon, p-value=0.991). (B) Emerging IJs from *S. littoralis* larvae cadaver for each strain. The amount of IJs exiting from larvae cadaver were not significantly different (Wilcoxon, p-value=0.31).

### Bacterial symbionts numeration in emerging IJs

For each strain, numeration of CFU in emerging IJs was performed after nematode crushing. This experiment revealed that after a cycle in the insect, several bacterial colonies displaying no luminescence appeared, indicating that they did not belong to the *Photorhabdus* genus. Therefore, only luminescent colonies were numerated. Results presented in Fig.4 show that there was slightly more *Photorhabdus* CFU numerated from nematode in symbiosis with the control strain (460+/-126 CFU) than with the *dam* overexpressing strain (270+/-100 CFU, p-value<0.01, glmm) (Fig 4). However, this experiment showed that each strain was able to colonize *H. bacteriophora*.

**Figure 4:**
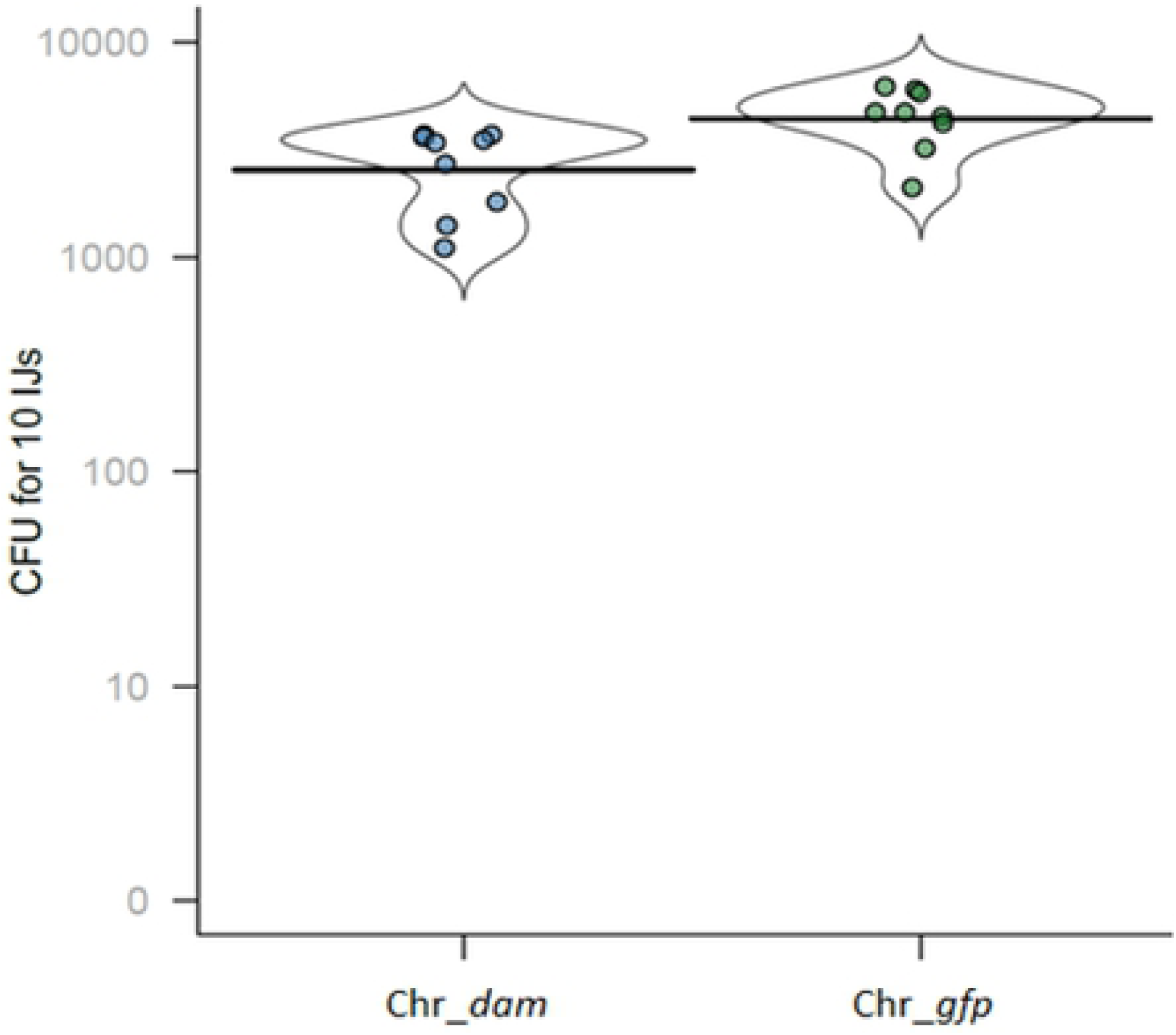
CFU in IJs nematodes for each strain. After crushing of 10 IJs and plating of the resulting suspension, CFU were numerated. A significant difference was observed between the two strains (glmm, p-value<0.01).

## Discussion

We previously described that Dam MTase allows the methylation of most (>99%) of the adenines in 5’-GATC-3’ motifs in the *P. luminescens* TT01 genome and that DNA methylation profile was stable during *in vitro* growth [43]. Dam DNA methylation is known to be involved in various phenotypes as pathogenicity in several bacteria as *S.* Typhimurium [19, 20], *Y. pestis* [22] or *A. hydrophila* [44]. The only studies about DNA methylation involvement in symbiosis are limited to bacterial-plant interactions: in *Bradyrhizobium* it is suspected to play a role in the cell differentiation to symbiotic stage [25] and in *Mesorhizobium loti* it is essential for nodulation [26, 27].

While we previously described that *dam* overexpression in *P. luminescens* causes a decrease in pathogenicity and motility [28], the role of Dam in the symbiotic stages of *P. luminescens* life-cycle remained to be investigated. Here, using a strain harboring an additional copy of the *dam* gene under the control of a constitutive promoter by a chromosomal insertion, we first confirmed that *dam* overexpression decreases motility and virulence in insect when compared to a control strain (Chr_*gfp*). The *in vitro* symbiosis between *H. bacteriophora* nematode and either the *P. luminescens dam*-overexpressing strain or the control strain showed similar amount of emerging IJs for each nemato-bacterial complex, revealing that the nematodes can feed and multiply on both strains *in vitro*. Three parameters were analyzed to determine the symbiosis efficiency of both strains *in vivo* after a cycle on insects: (i) The pathogenicity of the nemato-bacterial complex was assessed by recording the LT_50_, (ii) the nematode reproduction was assessed by numeration of IJs emerging from each cadaver, (iii) the bacterial ability to recolonize the nematodes gut inside the insect cadaver was assessed by numerating bacteria in IJs. The first two parameters (i.e. pathogenicity and emerging IJs) were done using two insect models in order to compare our results between a broadly used insect model (*G. mellonella*) and a more relevant insect for our nemato-bacterial complex (*S. littoralis*). Differences between the two insect models were observed. In *G. mellonella*, a significant difference of 2 hours in LT_50_ between both EPN complexes strains could be detected. In *S. littoralis*, a higher difference in LT_50_ was noted compared to that in *G. mellonella*, as a 6 hour-delay was required to kill half of the larval cohort for *Hb* Chr_*dam* strain compared to the control. No difference was observed in the number of emerging IJs between both EPN complexes after infestation of both insect models. Because in both insects the control strain took the same time to reach LT_50_ (48h) the observed difference between insect models is related to *dam* overexpression. One hypothesis is the involvement of Dam in genes regulation that are more important for the pathogenicity in *S. littoralis* model. Altogether these results show a decrease in pathogenicity of the nemato-bacterial complex overexpressing *dam* that can be caused, at least in part, by the decrease in pathogenicity of the bacteria alone, as previously described [28] and confirmed here. The observed differences in LT_50_ between injection and infestation with the two nemato-bacterial complexes in *S. littoralis* (2 hours delayed LT_50_ for Chr_*dam* strain by injection and 6 hours delayed LT_50_ for *Hb* Chr_*dam* by infestation) suggest a role of Dam not only in the bacterial pathogenicity, but also in the pathogenicity of the nemato-bacterial complexes. Because a longer time is required for the nemato-bacterial complexes to kill insects than for the bacteria alone (48h vs 36h, respectively for the control strain), another hypothesis might be that this difference is only a knock-on effect. Here, we show that both tested bacterial strains allow nematode multiplication *in vitro* and *in vivo*, nematode virulence in insects, nematode emergence from the cadavers, and nematode’s gut colonization, revealing that symbiosis establishment is not impaired by the bacterial *dam* overexpression. However, we cannot rule out that the observed slight reduction in the amount and CFU per IJ can play a role in life history trait of the nemato-bacterial complex. This could be investigated in further studies by monitoring the evolution of the three parameters analyzed here (pathogenicity, emerging IJ, amount and CFU per IJ) after several successive cycles of infestation.

## Conclusion

This study showed that the *P. luminescens* Dam contribute to the pathogenicity in *S. littoralis* after injection of the bacteria alone and to a greater extent after infestation by the nemato-bacterial complex. However, overexpression of the *P. luminescens dam* gene does not significantly play a role in the symbiotic stages with the nematode.

## Author contributions

AP performed the experiments and analyzed the data; AP, DB, AL, DC, AG, JB designed the experiments; AP, SP, MC performed statistical analyze; AP wrote the manuscript. DC, AG, JB critically revised the manuscript. All authors read and approved the final version of the manuscript.

## Acknowledgments

The authors thank the quarantine insect platform (PIQ), member of the Vectopole Sud network, for providing the infrastructure needed for pest insect experimentations.

## Supporting Information

**Table S1:** Primers used in this study

**Table S2:** Phenotypes of *P. luminescens* TT01 Chr_*dam* and Chr_*gfp* strains

**Figure S1**: Emerging IJs from *in vitro* symbiosis association

